## Supplementary information for "STREAMLINED PROTEOME-WIDE IDENTIFICATION OF DRUG TARGETS INDICATES ORGAN-SPECIFIC ENGAGEMENT"

SUPPLEMENTARY INFORMATION FOR STREAMLINED IDENTIFICATION OF DRUG TARGETS INDICATES ORGAN-SPECIFIC ENGAGEMENT

Figure S1: TPP melting point distributions

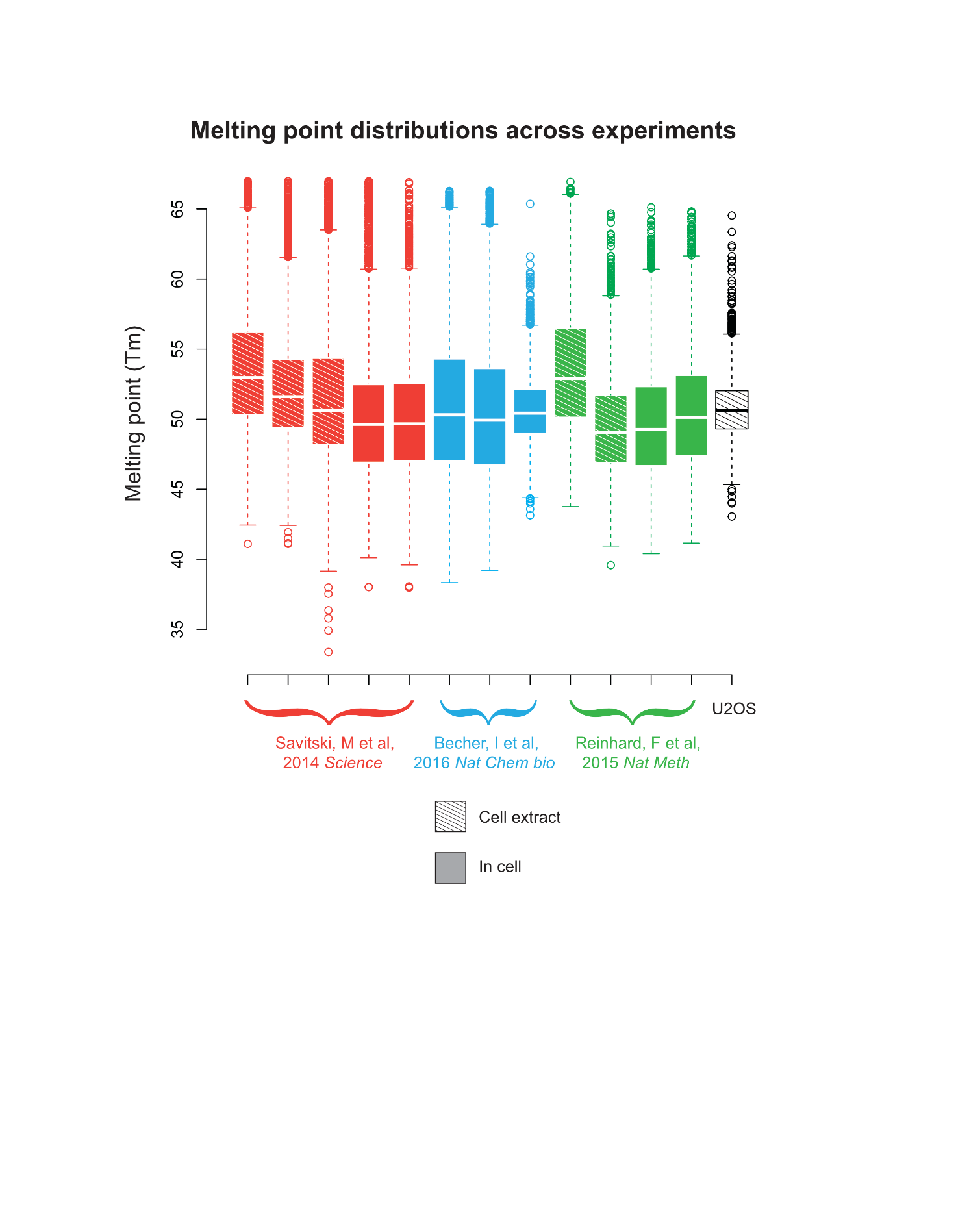

Figure S1: Published melting point box plot distributions from different Thermal Proteome Profiling (TPP) experiments^1–3^ are plotted including from U2OS generated from this study.

Figure S2: Comparison between DIA and TMT for TPP-PISA experiments

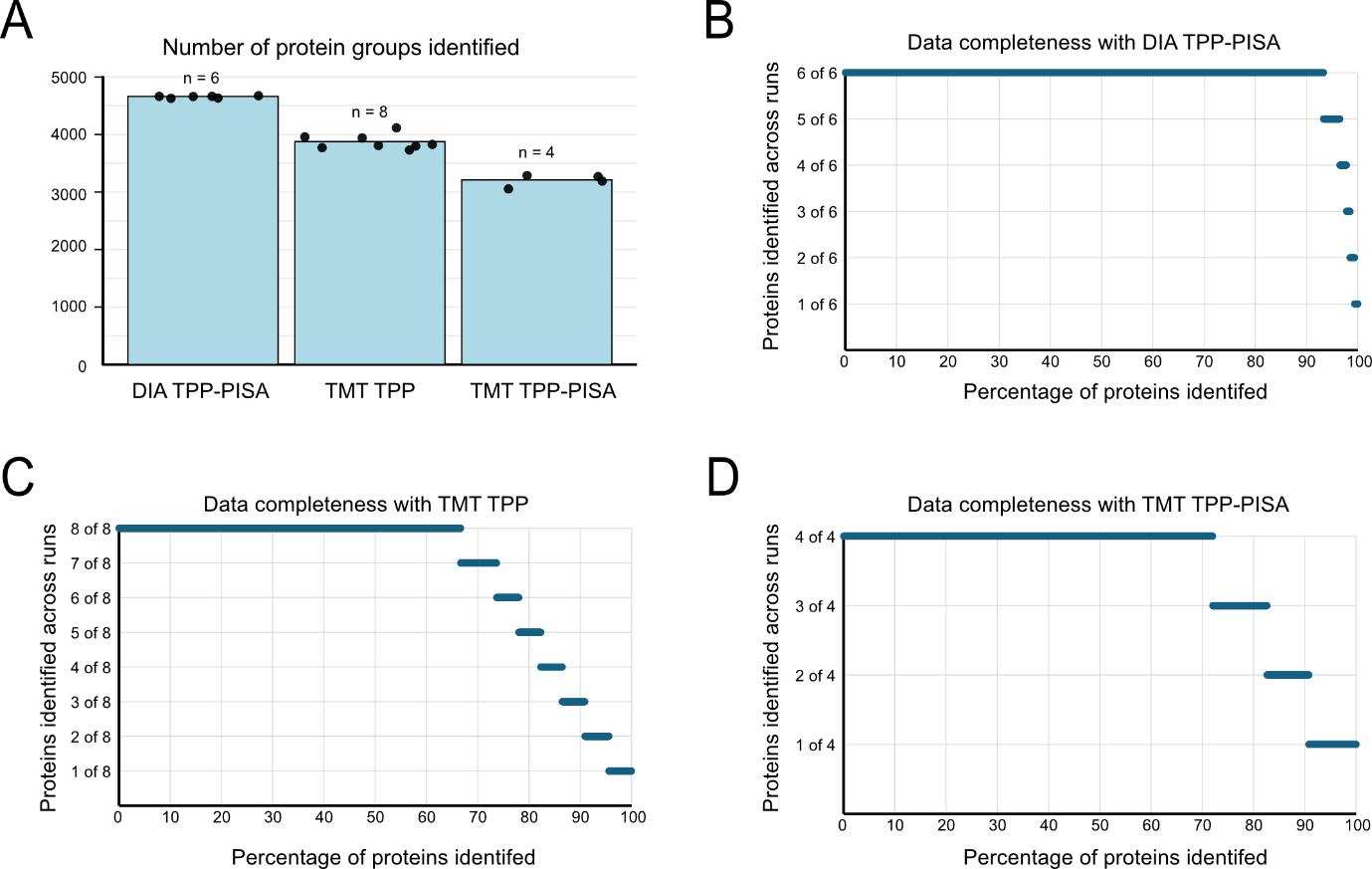

Figure S2: Evaluation of protein identification rates and data completeness for TPP-PISA experiments using TMT labeling approach (DDA) vs DIA mass spectrometry analysis. A) Number of proteins identified across different experiments. Each data point represents an experiment with the number of identified proteins (protein groups). B) Data completeness analysis of DIA TPP-PISA for the DMSO (n=3) and Staurosporine (n=3) experiment used for this analysis. X-axis displays percentage of data which was identified in the different replicates, i.e. >90% of the proteins were identified in 6 of 6 runs (y-axis) for this experiment. Similar data completeness analysis for C) TMT TPP and D) TMT TPP-PISA.

Figure S3: 96-well strategy for TPP-PISA experiments

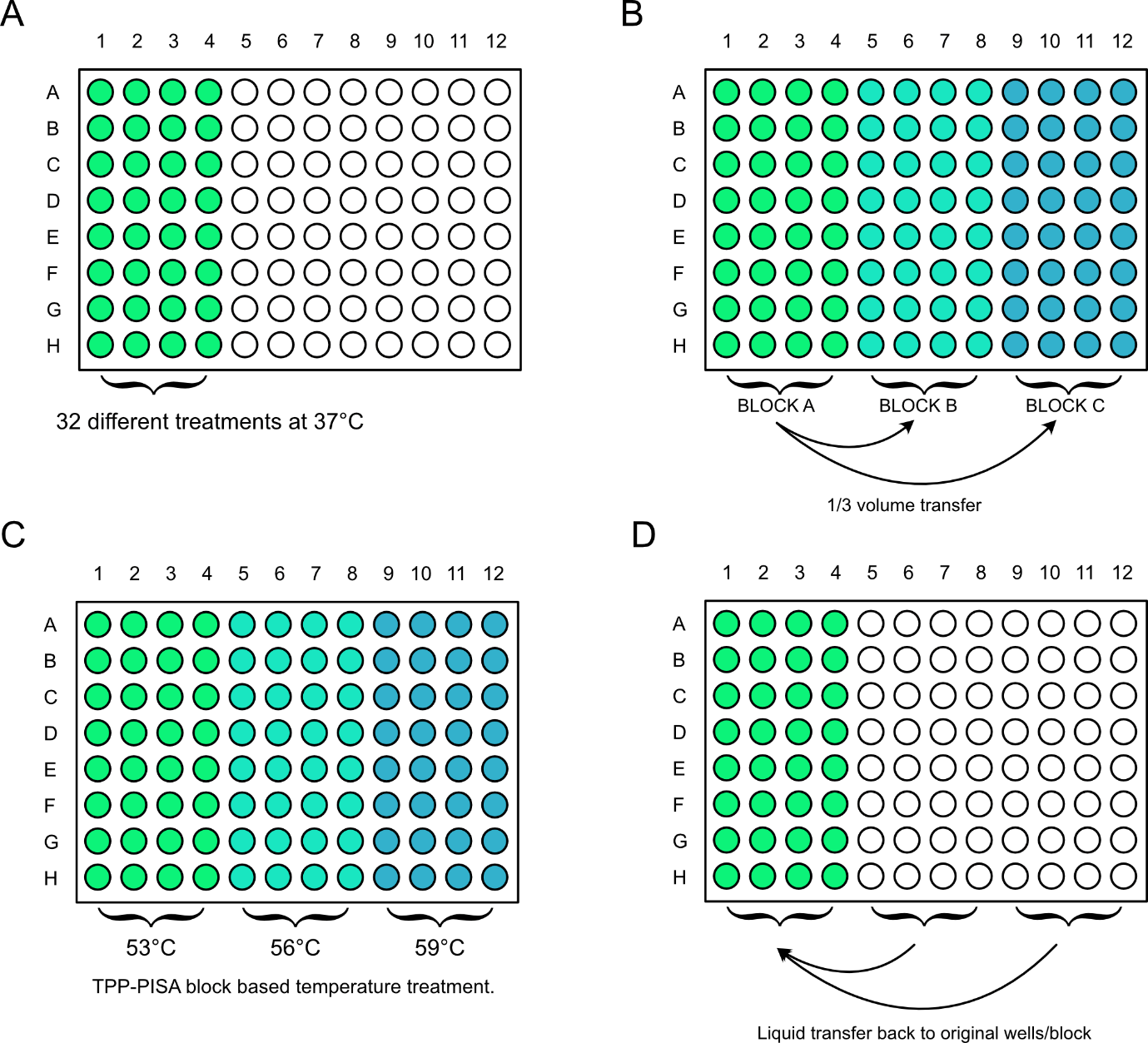

Figure S3: The strategy for 96-well format for TPP-PISA experiments which enables up to 32 different treatments in one plate. A) Protein extracts can be aliquoted into rows A-H and columns 1-4 in PCR plates followed by the addition of various drugs and ligands. The mixture is then heated at 37⁰C for 10 minutes. B) After experimental treatment, equal volume is distributed to three blocks. This is achieved through transfer of 1/3 of the volume from block A to block B and C, resulting in equal volume in all wells and blocks. C) The blocks are heated at 3 different temperatures as described for the TPP-PISA experiments. D) The liquid volume is transferred back to the original wells and block (block A) from block B and C after temperature treatment.

Figure S4: Staurosporine response in Rat organ extracts

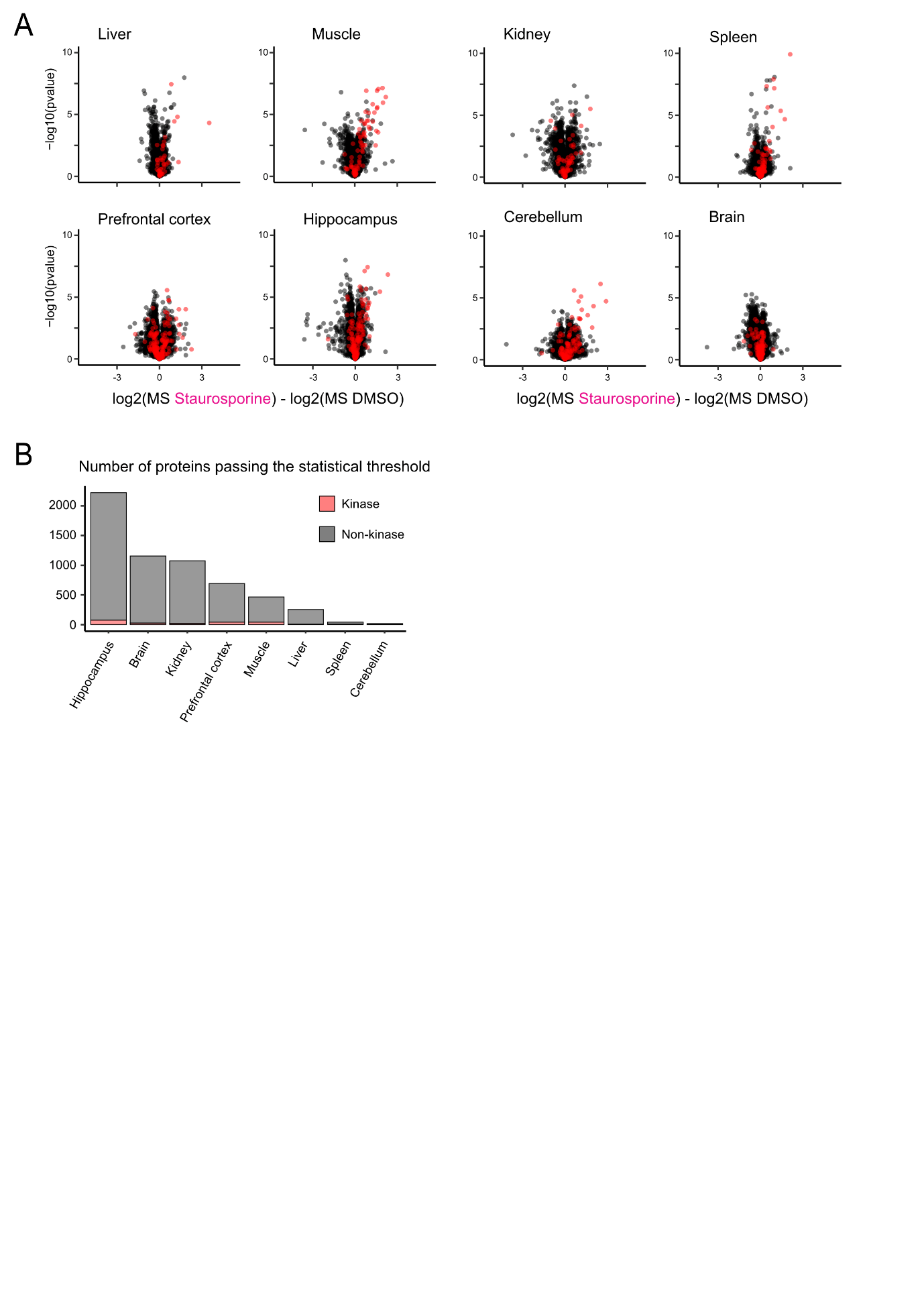

Figure S4. A) Volcano plots for all the statistical analysis of rat organs subjected to TPP-PISA with the protocol presented in (A). -log10(p-values) are plotted on the vertical axis, and the differences between log2-transformed protein groups quantities in the soluble fractions of the staurosporine- and DMSO-treated conditions are presented on the horizontal axis. Kinases are highlighted in red. B) Number of hits found in different rat organ extracts with q-value <0.05.

Figure S5: Heatmap of Staurosporine targets in Rat organ extracts

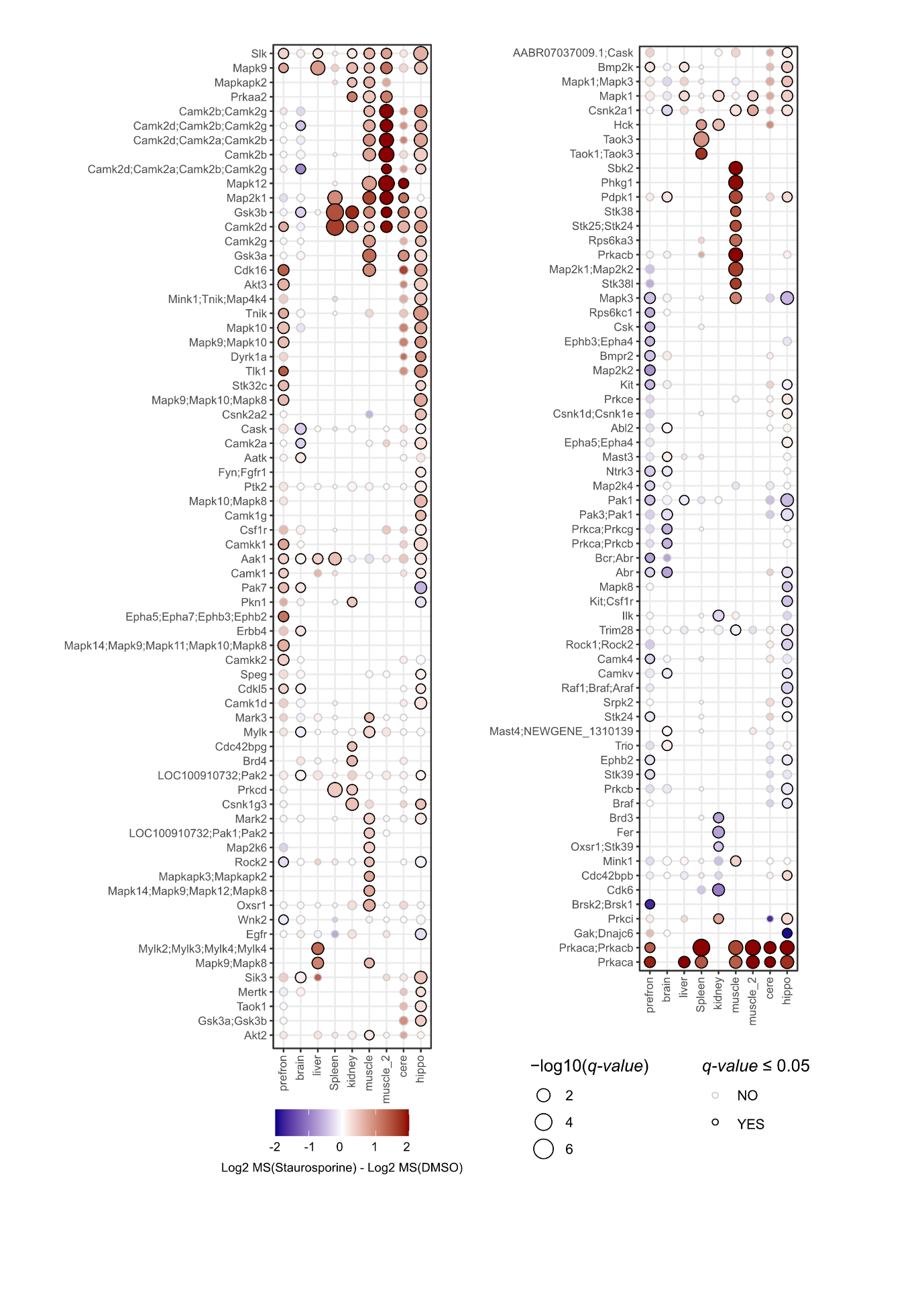

Heatmap of all Staurosprorine kinase targets observed in rat organ extracts. Normalized fold change values log2(MS Staurosporine) – log2(MS DMSO) are color coded for each point, and the size of the point reflects the *q-value*. Stroke color for each points displays whether the kinases pass statistical threshold based on fold change and *q-value*.

Figure S6: Validation of hPirin positive control inhibitor TPhA by SPR

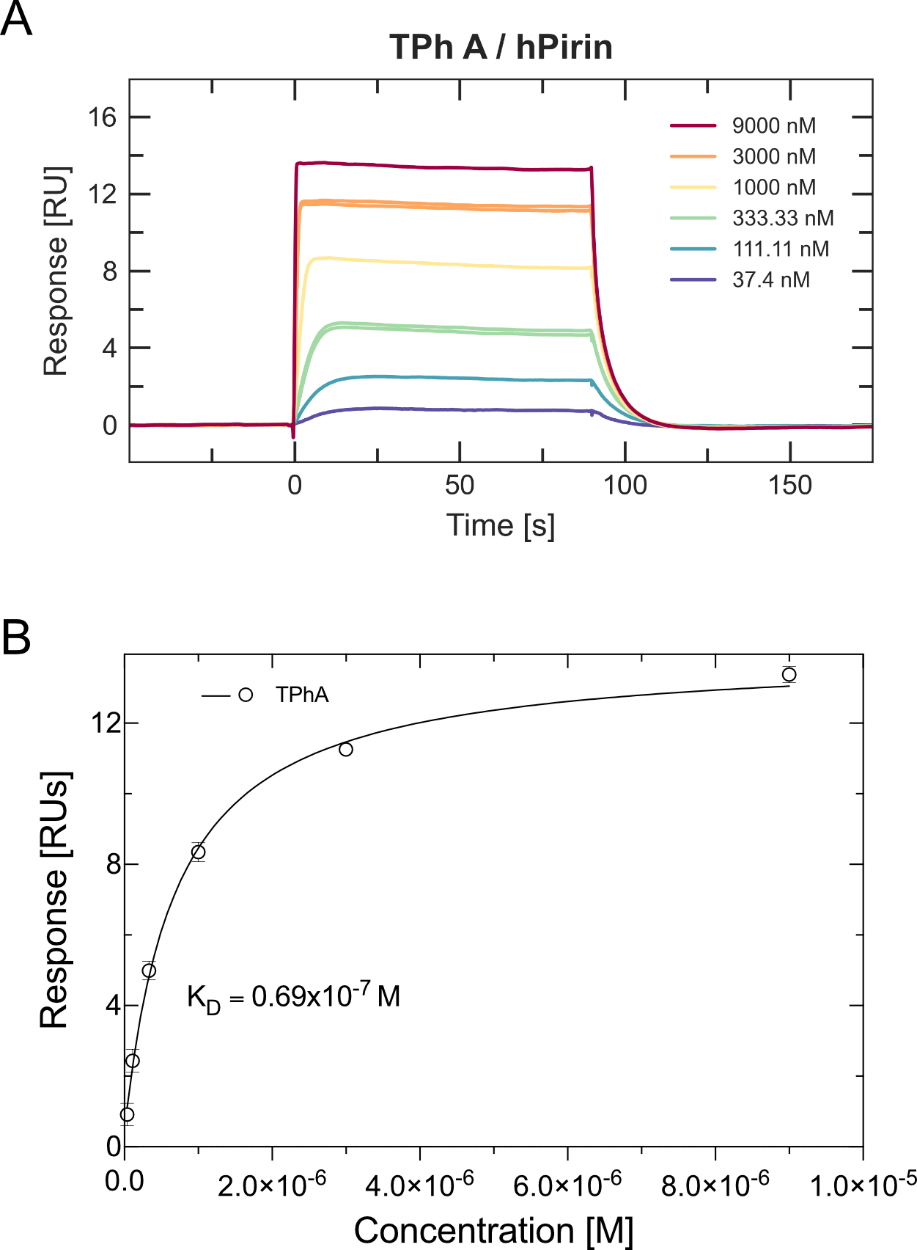

Figure S6. **A)** SPR sensorgram of recombinant human Pirin (hPirin) and Triphenyl Compound A (TPhA) showing a dose dependent response. **B)** Equilibrium dissociation constant (Kd) determined from the steady state model of TPhA for hPirin confirming its high specificity.

Figure S7: HexB assay in HeLa protein extracts

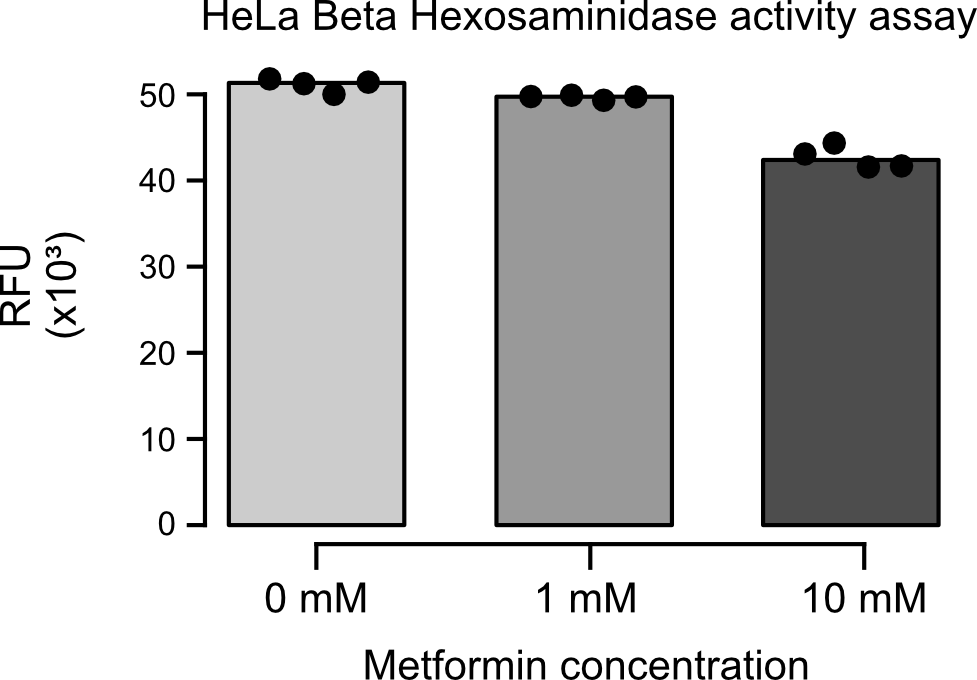

Figure S7. HexB activity performed on HeLa cellular extracts using a fluorescent assay with 1mm and 10mm metformin. Relative fluorescence units are plotted. Individual replicates are plotted (n=4).
